# Single particle profiler for measuring content and properties of nano-sized bioparticles

**DOI:** 10.1101/2022.07.08.499323

**Authors:** Taras Sych, Jan Schlegel, Hanna M.G. Barriga, Miina Ojansivu, Leo Hanke, Florian Weber, R. Beklem Bostancioglu, Kariem Ezzat, Herbert Stangl, Birgit Plochberger, Jurga Laurencikiene, Samir El Andaloussi, Daniel Fürth, Molly M. Stevens, Erdinc Sezgin

## Abstract

It is technically challenging to study the content and properties of nanoscale bioparticles in a high-throughput and single-molecule manner. We developed a high-throughput analysis method, called single particle profiler (SPP) that provides single-particle information on content and biophysical properties of thousands of particles. We applied SPP to measure the mRNA encapsulation efficiency of lipid nanoparticles, viral binding efficiency of different nanobodies and biophysical heterogeneity of liposomes, lipoproteins, exosomes and viruses.

## Main text

Physiological nanometer-sized particles in the human body are of utmost importance for health and disease. For example, lipoproteins (5 nm (HDL) – 80 nm (VLDL), vesicular monolayers of lipids and proteins encapsulating cholesterol esters and triglycerides) transport lipids to maintain cellular metabolism^1^; extracellular vesicles (EVs, vesicular bilayers encapsulating various biomolecules including nucleic acids and proteins) (< 200 nm) take part in immune responses, cell-cell-communication and signaling^2^; and viruses with an average size of 100 - 200 nm cause a variety of diseases. Moreover, synthetic liposomes (vesicular bilayers) and lipid nanoparticles (LNPs, vesicular monolayers) are widely employed for drug delivery and vaccines^3^. The analysis of their content and biophysical properties can shed light on their structure, function, and behavior in health and disease. Most of the existing methods to study bioparticles rely on biochemical analysis, mass spectroscopy, total internal reflection fluorescence (TIRF) microscopy and conventional flow cytometry. Biochemical methods and mass spectroscopy are bulk methods (i.e., lack single particle sensitivity). TIRF microscopy provides excellent signal to noise ratio in truly single particle manner; however, it requires fixation of the vesicles on the surface and yields low-throughput data. Flow cytometry, on the other hand, is a high throughput method; however, it is generally suitable for cell-sized objects. Recently, there have been attempts to analyze small EVs with flow cytometry^4–9^ and there is an ongoing development of the “nano-flow” devices that rely on microfluidic equipment^10,11^. However, these methodologies are still limited to average particle size of >200 nm and require dedicated and often costly equipment.

We designed a method, single particle profiling (SPP), that is based on analysis of fluorescence fluctuations of thousands of diffusing particles in solution, recorded with a commercially available confocal microscope (Figure 1a). Briefly, fluorescently labeled particles diffuse through the diffraction-limited observation volume (≈0.5 fL) where the fluorescence emission from multiple channels is monitored continuously (as in fluorescence cross-correlation spectroscopy (FCCS)^12,13^). Unlike the FC(C)S which reduces all fluctuations into a single curve^14^, we identify individual peaks in intensity fluctuations in multiple channels using the custom, freely available python script (Figure 1b, see Supporting Movie 1 for the tutorial of the software and Github for downloading the software). Based on the intensity of each individual peak in multi-channels, we can construct a density plot (Figure 1c) and histograms for each individual channel (Figure 1d). Therefore, like flow cytometry, this approach can be used to measure the fluorescence intensities in single nanometer-sized particles smaller than 250 nm. Such information can be used for *content measurement* and *biophysical profiling* of particles.

**Figure 1.**
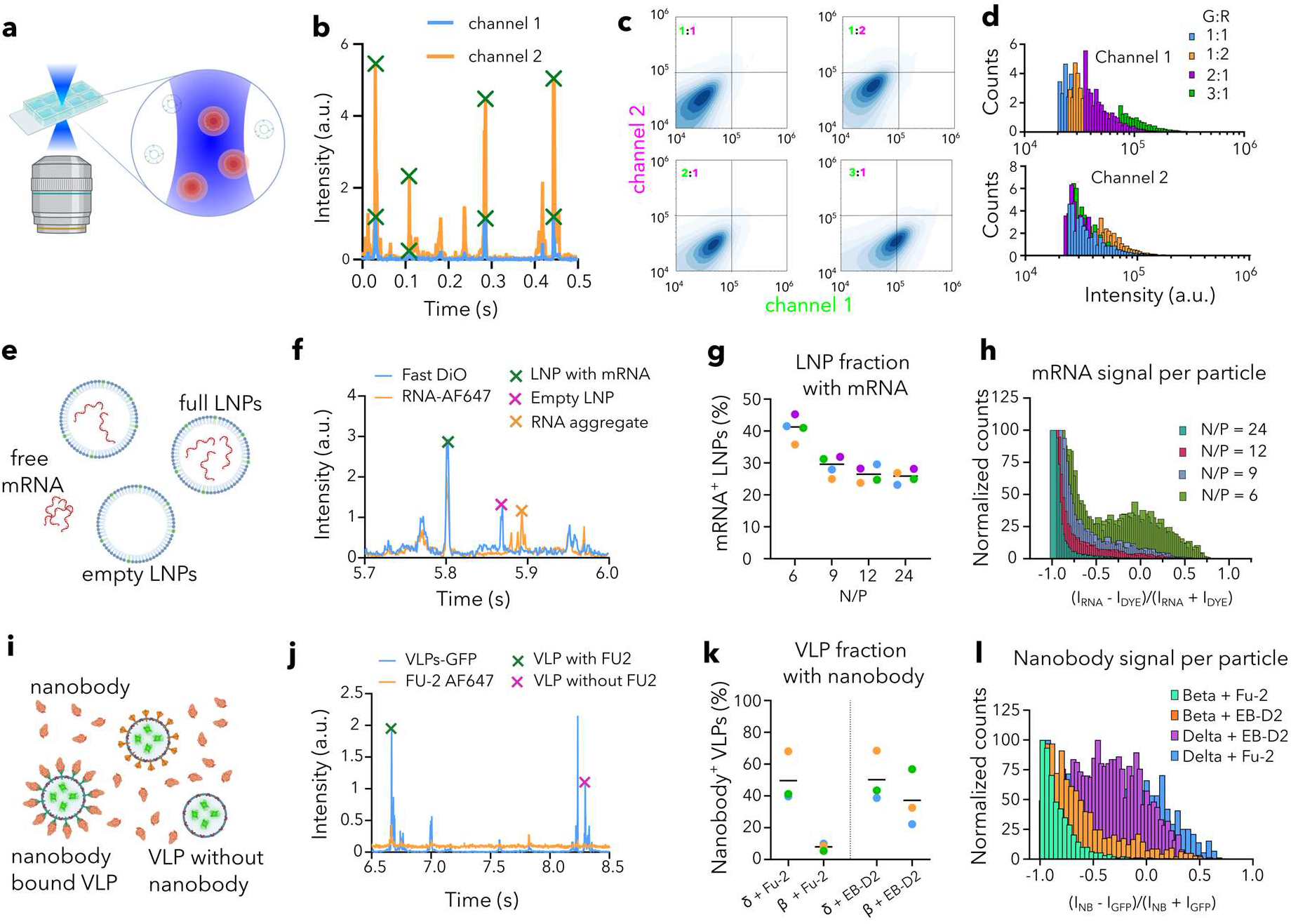
Single Particle Profiling for content analysis. **a)** SPP setup; **b)** Fluorescence intensity traces and peak calling; **c)** 2D density map of fluorescence intensities for liposomes loaded with Fast-DiO and Abberior Star Red DPPE at different ratios; **d)** Intensity distributions for each channel for the samples shown in panel c; **e)** Scheme for mRNA encapsulation of LNPs; **f)** Representative peaks for possible scenarios; **g)** LNP fraction with mRNA signal for different N/P ratios (colors represent the mean of one replicate); **h)** Ratiometric histogram of mRNA vs lipid dye intensity for single LNPs; **i)** Scheme for nanobody binding to VLPs; **j)** Representative peaks for possible scenarios; **k)** VLP fraction with nanobody signal for different nanobodies and different variants (colors represent the mean of one replicate); **l)** Ratiometric histogram of nanobody vs lipid dye intensity for single VLPs.

First, to demonstrate the ability of SPP for *content profiling*, we prepared liposomes loaded with either green molecules and/or red molecules incorporated into the lipid membrane. Single- or double-loaded (only green dye, only red, or both) confirmed the applicability of our methodology for content profiling (Supplementary Figure 1). To test the content differences accessible by SPP, we changed the ratio of green (Fast-DiO) and red (Abberior Star Red DPPE) signal in liposomes (1:1, 1:2, 2:1 and 1:3) which was successfully captured by SPP (Figure 1c, d). To show the high throughput capacity of SPP, we performed this experiment in multi-well plates (Supplementary Figure 2).

In SPP measurements, each peak is analyzed for its brightness, width, and co-occurrence with another color (Supplementary Figure 3). Peak brightness can be used for determining the clustering of particles (Supplementary Figure 4) and combined with peak width analysis, it can detect the presence of small number of large particles or aggregates (Supplementary Figure 5). Furthermore, co-occurrence of the same peak in multiple channels and analysis of corresponding intensities provide information on particle content. Conventional co-diffusion analysis fails to evaluate co-occurrence robustly in heterogenous samples because a single bright peak (which represents a large-sized impurity in the sample) skews the cross-correlation analysis (Supplementary Figure 6). To use this advantage of SPP, we applied it to study two important biological challenges: measuring *i)* mRNA encapsulation efficiency of LNPs; and *ii)* antibody binding to virus particles.

LNPs are widely used as drug delivery and vaccine delivery agents^15^. One of the crucial parameters for effectiveness of LNP-based treatments is the payload encapsulation efficiency. We applied SPP to measure the encapsulation efficiency for LNPs loaded with mRNA (Figure 1e). Current employed measurements for mRNA encapsulation efficiency of LNPs are based-on bulk measurements using RNA binding dyes such as RiboGreen (Supplementary Figure 7). This assay measures the fluorescence signal from the RiboGreen dye in the sample before vs. after detergent treatment which are, respectively, proportional to non-encapsulated mRNA in solution vs. total mRNA. This ratio is termed as the “encapsulation efficiency” and is measuring the percentage of mRNA encapsulated in LNPs at the population level, but does not yield insights on the empty vs full LNPs. As such, even if the RiboGreen yields 100% encapsulation efficiency (i.e., all mRNA in the sample is inside LNPs), it is still possible that only a fraction of the LNPs carry all the cargo, while another pool is empty. Using SPP, however, we directly visualized and quantified the encapsulation by measuring the mRNA and lipid-dye signal co-occurrence of each particle (Figure 1f). Control LNPs with two lipid dyes showed ≈100% co-occurrence, confirming the robustness of SPP for such measurements (Supplementary Figure 7). For encapsulation analysis, each LNP (blue peaks in Figure 1f) was quantitatively analysed for the mRNA content (i.e., whether it contains mRNA and how much). We calculated the fraction of LNPs showing mRNA signal, i.e., full LNP fraction (Figure 1g), and importantly, encapsulation heterogeneity, i.e., how much mRNA each LNP has (Figure 1h, “-1” corresponds to empty LNPs and higher values mean more mRNA inside LNPs, see Methods for calculations). By changing the charge ratio of ionizable lipid:mRNA (N/P ratio), we showed that most efficient encapsulation (both fraction of full LNPs and amount of mRNA inside each LNP) was at N/P=6. This shows that LNP formulation parameters affect both the cargo encapsulation and the extent of amount loaded per LNP, which have crucial implications on the performance of LNPs as drug carriers. Unlike the currently prevailing bulk methods such as RiboGreen assay, which can only give an overall percentage for the loading efficiency without any insight to the cargo distribution, SPP provides highly valuable single-particle information for the development and quality control of LNP-based drug formulations.

Virus neutralization by antibodies is key for immunity. It is essential to swiftly determine whether certain antibodies bind the strain of interest. We applied SPP to evaluate the binding efficiency of nanobodies specifically developed to bind SARS-CoV-2 Spike protein. To this end, we generated different variants of SARS-CoV-2 virus-like particles (beta and delta) carrying GFP and incubated them with nanobodies Fu2 and EB-D2 (that target the receptor binding domain (RBD), and an epitope outside the RBD, respectively) site-specifically conjugated to a fluorescent dye^16–18^ (Figure 1i). SPP analysis shows virus peaks that do or do not co-occur with nanobody peaks (Figure 1j). Quantification of the co-occurrence shows that ≈50% of Delta and only ≈10% of Beta VLPs were bound by Fu2. On the other hand, ≈50% of Delta and ≈40% of Beta VLPs were bound by the EB-D2 nanobody (Figure 1k). Normalized intensity histograms showed how much antibody bound to each virus particle: Beta/Fu2<Beta/EB-D2< Delta/EB-D2< Delta/Fu2 (Figure 1l). Thus, SPP is sensitive to detect binding differences between nanobodies to different viral variants paving the way to antibody neutralization screening.

Co-occurrence of particle fluorescence signals in multiple channels can also be used for studying biophysical properties of nanosized bioparticles using ratiometric environment sensitive probes (Supplementary Figure 8). To demonstrate the ability of SPP for such *biophysical profiling*, we prepared liposomes of distinct lipid compositions and supplemented them with 0.1 mol % of the ratiometric dye Nile Red 12 S (NR12S, Supplementary Figure 1). The emission spectrum of this dye is sensitive to fluidity of the lipid environment which is measured with an index called Generalized Polarization (GP). Numerical value of GP changes between +1 and -1 and is inversely proportional to membrane fluidity (e.g., higher GP = lower fluidity). We prepared liposomes of different membrane fluidity by using saturated lipids, unsaturated lipids (18:1/18:1 DOPC, 18:1/16:0 POPC, 16:0/16:0 DPPC) and cholesterol in various combinations. As expected, membrane fluidity increased together with lipid saturation degree and cholesterol content (Figure 2a, Supplementary Figures 9). Importantly, single particle capability of SPP allows us to perform advanced statistical analysis of GP data and histograms. For example, sigma (σ) of the GP histogram provides information on the heterogeneity of the sample, that is, smaller σ means more homogeneity. This was demonstrated for the liposome mixtures (inset graphs in Figure 2a, Supplementary Figures 9). The biophysical heterogeneity of the liposomes increased with the complexity of the lipid mixture. A key advantage of SPP compared to bulk techniques is the ability to extract single particle readings from thousands of particles, which allows us to dissect multicomponent mixtures. To show this, we prepared a mixture consisting of POPC liposomes and DPPC/chol liposomes. Analysis of such mixture indeed revealed the presence of two liposome populations: one population of low order (POPC liposomes) and a second population of high order (DPPC/chol liposomes) (Figure 2b). Moreover, even when the mixed populations are similar in their biophysical properties and cannot be easily separated using multi-component analysis, one can still infer on the heterogeneity by measuring σ of the GP distribution (Supplementary Figure 10).

**Figure 2.**
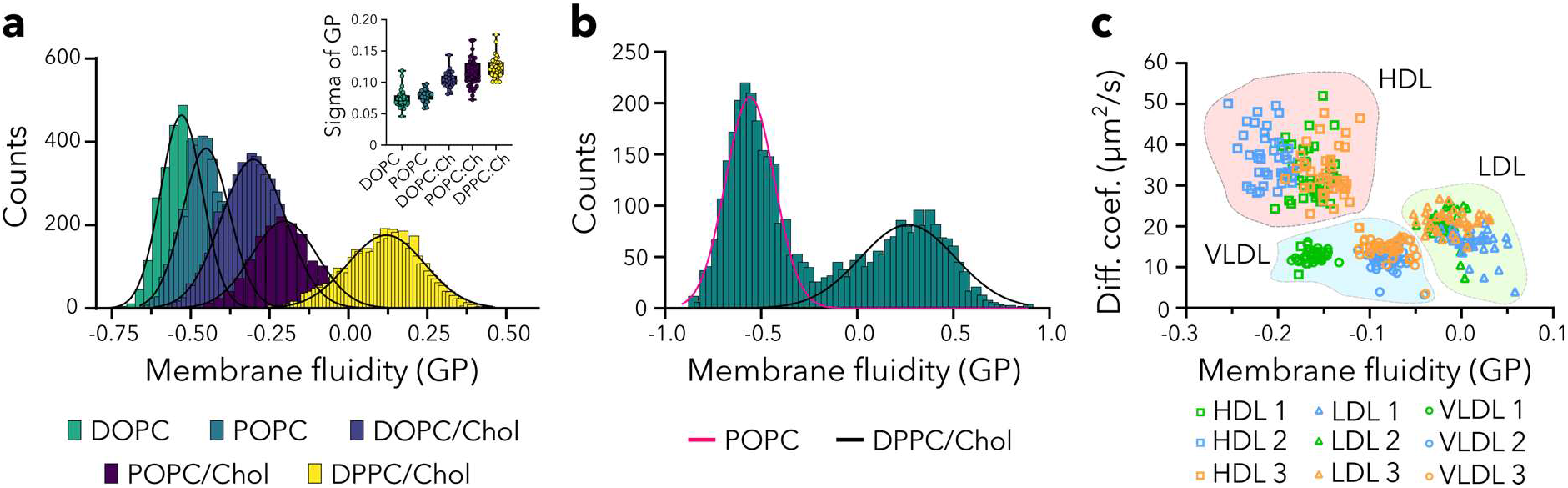
Biophysical properties of nanosized bioparticles by SPP. **a)** The histograms of generalized polarization for liposomes of different lipid compositions and inset for the σ of GP distribution; **b)** The GP histogram of the mixture of liposomes with distinct composition shows two distinct populations; **c)** Diffusion coefficient vs. GP dot plot for lipoproteins from three different donors.

One of the unique advantages of SPP compared to flow cytometry tools is the ability to measure diffusion of particles since it is based on fluorescence fluctuations. Two parameters instead of one generally provide better separation power (Supplementary Figure 11). Here, diffusion of particles (directly related to particle size, Supplementary Figure 12) can be an additional parameter to separate different particles. To demonstrate this, we used blood plasma from healthy donors and isolated three major types of lipoproteins (LPs): high-density lipoprotein (HDL), low-density lipoprotein (LDL) and very low-density lipoprotein (VLDL). LP heterogeneity is a critical factor for cholesterol homeostasis and several disorders including cardiovascular diseases^19^. We labelled all LPs with NR12S and performed SPP and multi-parameter analysis. Separation of LDL particles from VLDL particles was not clear using either diffusion or GP (i.e., notable overlap for both parameters), but combining these parameters led to a cleaner separation (Figure 2c, Supplementary Figure 13). Moreover, donor-to-donor differences were also evident from these analyses. We performed similar analysis for liposomes, EVs, virus-like particles and lipid nanoparticles (Supplementary Figure 14-18). These analyses show that SPP can provide multi-parametric information (e.g., biophysical properties and diffusion of particles) and such multi-parametric separation of nanosized bioparticles can be used in the context of health and disease.

Here, we presented and validated a new analysis method for high throughput single particle profiling. We showed that *(i)* content and biophysical properties of nanosized bioparticles can be studied with SPP in single-particle and high-throughput manner; *(ii)* sample heterogeneity can be studied using statistical analysis and *(iii)* multiple parameters (such as diffusion and fluidity) can be obtained for clustering analysis. We used these features of SPP to study mRNA encapsulation efficiency for LNPs, viral binding efficiency of nanobodies and biophysical heterogeneity of nanosized bioparticles. This method is based on commercially available and highly accessible confocal systems and has wide applicability for resolving content and organization of the nanosized physiological particles. SPP does not require dedicated equipment and is not limited to fixed spectral regions thanks to the available detector flexibility in confocal systems. SPP provides information on lipid and protein content, biophysical parameters and polydispersity of nano-sized bioparticles, which can shed light on the role of these particles in health and diseases.

## Data availability statement

All raw data will be available at the service FigShare upon publication (doi: 10.17044/scilifelab.20338869).

## Code Availability

The source code of the script as well as the standalone distributions are available at github: https://github.com/taras-sych/Single-particle-profiler.

## Acknowledgements

This work is supported by National Microscopy Infrastructure Sweden (VR-RFI 2016-00968), SciLifeLab National COVID-19 Research Program financed by the Knut and Alice Wallenberg Foundation, Karolinska Institutet, SciLifeLab, Swedish Research Council Starting Grant (2020-02682) and Human Frontier Science Program (RGP0025/2022).

## Author contributions

T.S., J.S., H.M.G.B., M.O., L.H., F.W., R.B.B., K.E., H.S., B.P., J.L. and D.F. prepared samples, T.S., J.S., H.M.G.B., M.O. and ES performed the experiments, S.E.A., M.M.S., E.S. supervised the students, T.S. wrote the code, T.S. and E.S wrote the manuscript, all authors contributed to the manuscript writing.

