## Supplementary Information for "Single particle profiler for measuring content and properties of nano-sized bioparticles"

### Materials and Methods

**Preparation of LUVs.** 1,2-Dioleoyl-sn-glycero-3-phosphocholine (DOPC), 1-palmitoyl-2-oleoyl-sn-glycero-3-phosphocholine (POPC), Dipalmitoylphosphatidylcholine (DPPC) and cholesterol were from Avanti polar lipids. Fluorescent lipids and lipid-like probes we used: Abberior Star Red DPPE (Abberior), TopFluor Cholesterol (Avanti), FastDiO (ThermoFisher), NR12S was provided by Dr. Andrey Klymchenko, University of Strasbourg. Five lipid mixtures in chloroform at 1 mg/ml were prepared: pure DOPC, pure POPC, DOPC/chol (70/30), POPC/chol(70/30), DPPC/chol(70/30). Additionally, lipid mixtures with differen cholesterol percentage were prepared: POPC/chol(95/5), POPC/chol(90/10), POPC/chol(85/15), POPC/chol(80/20), POPC/chol(75/25), POPC/chol(70/30), POPC/chol(65/35), POPC/chol(60/40), POPC/chol(55/45), POPC/chol(50/50). Finally, mixtures containing POPC with different ratios of fluorescent lipids were prepared: POPC/FastDiO/Abberior Star Red PE (99.8/0.1/0.1), POPC/FastDiO/Abberior Star Red PE (99.7/0.1/0.2), POPC/FastDiO/Abberior Star Red PE (99.7/0.2/0.1), POPC/FastDiO/Abberior Star Red PE (99.6/0.3/0.1), POPC/TopFluor Cholesterol/Abberior Star Red PE (99.8/0.1/0.1), POPC/TopFluor Cholesterol (99.9/0.1), POPC/ Abberior Star Red PE (99.9/0.1). Mixtures were dried under the flow of nitrogen, rehydrated with buffer (150 mM NaCl, 10 mM Hepes, 2 mM $CaCl_2$) and vortexed harshly to form multilamellar vesicles. Then, the suspension of MLVs was sonicate at power 3, duty cycle 40% for 10 mins using Branson Sonifier 250. The size of the resulting vesicles was checked using dynamic light scattering (Malvern Zeta-sizer). LUVs were stored at 4 degrees under nitrogen and the solutions of LUVs (1 mg/ml) were incubated with Nile Red 12 S (NR12S, 1 μM in DMSO) directly before profiling.

**Isolation of lipoproteins.** Blood plasma was obtained from Blood transfusion station of Karolinska Hospital, Stockholm, donated by apparently healthy normolipidemic volunteers. Lipoprotein particles were isolated as previously described via sequential flotation ultracentrifugation[1]. Briefly, the density of blood plasma was sequentially adjusted using KBr to 1.019 g/l, 1.063 g/l and 1.22 g/l in order to isolate very low density lipoprotein (VLDL), low density lipoprotein (LDL) and high density lipoprotein (HDL) respectively. Lipoproteins were stored at 4 degrees under nitrogen and the solutions of LPs (1 mg/ml in PBS) were incubated with Nile Red 12 S (NR12S, 1 μM in DMSO) directly before profiling.

**Preparation of fluorescent RNA:** Fluorescently labeled RNA was synthesized by in vitro transcription (E2040S, NEB Inc.). One microgram of dsDNA template containing the T7 RNA promotor upstream from the Firefly luciferase gene (FLuc, 1.8 kb length, N0426, NEB Inc.) was added together with NTPs, 0.75X T7 RNA polymerase buffer, and 75 units of T7 RNA polymerase in a 20 μl reaction volume. The reaction was incubated overnight at 37°C. NTPs were at a final concentration of 7.5 mM except for UTP, which had a final concentration of 5 mM and was supplemented with 1.24 mM 5-Ethynyl-UTP (CLK-T08, Jena Biosciences Gmb). The addition of 5-Ethynyl-UTP results in alkyne-functionalized RNA, which was subsequently processed via Cu(I)-catalyzed (azide-alkyne) cycloaddition (CuAAC) to add fluorophores onto the RNA. After in vitro transcription, RNA was purified by silica spin column (R1055, Zymo Research Inc.). Fluorophores (AZDye 647 and AZDye 488) were added to the RNA by CuAAC using fluorophores conjugated to azides with a copper-chelating system in their structure (1475 and 1482, Click Chemistry Tools Inc.). Ligand BTTAA (50 mM, CLK-067, Jena Biosciences ) was allowed to react with 10 mM of CuSO4 (v800132, Sigma Aldrich) in nuclease-free water. Meanwhile, the functionalized RNA was eluted in 22.2 µl of nuclease-free water and supplemented with 1.8 µl of 10 mM azide-fluorophore dissolved in DMSO (276855, Sigma Aldrich). 60 mg of sodium L-ascorbate was dissolved in 1 mL of nuclease-free water. Immediately after the ascorbate was fully dissolved, it was added to the BTTAA/CuSO4 mixture at a final concentration of 100 mM. Six microliters of the BTTAA/CuSO4/ascorbate reaction were then added to the azide-fluorophore and RNA reaction, which was degassed using argon and sealed with parafilm, and incubated overnight. Labeled RNA was purified by silica spin column (R1055, Zymo Research Inc.) and measured on NanoDrop (ThermiFisher Inc.). The fluorescently labeled RNA was ethanol precipitated overnight using 3M sodium acetate to create a pellet for further experiments.

**Preparation of lipid nanoparticles.** 1-Oleoyl-rac-glycerol (monoolein) was purchased from Sigma Aldrich (Merck) and cholesterol (ovine wool), 1,2-dioleoyl-3-trimethylammonium-propane (18:TAP; DOTAP) and 1,2-dimyristoyl-sn-glycero-3-phosphoethanolamine-N-[methoxy(polyethylene glycol)-2000] (14:0 PEG2000 PE; DMPE-PEG2000) from Avanti Polar Lipids. All components were used without further purification. To make lipid films, the lipids were dissolved in chloroform (10 – 20 mg / mL) and mixed in the appropriate volumes to obtain the desired molar ratios (monoolein : cholesterol : DOTAP : DMPE-PEG2000 60 : 30 : 10 : 2.5 mol%). Lipid films were further supplemented with 0.5 mol-% Fast DiO dye. Chloroform was evaporated at room temperature overnight. Lipid films were subsequently stored under nitrogen, sealed at -20°C until LNP preparation. For LNP preparation, lipid films were defrosted and dissolved in absolute ethanol in concentration of 16.7 mM. Cargo was dissolved in 0.025 M pH 4 sodium acetate buffer in concentrations giving N/P ratios of 0, 6, 9, 12 and 24 in the final LNP formulations. LNPs were formulated using a syringe pump (The Harvard Apparatus Pump 33 dual drive system). Both solutions were loaded in 2.5 mL Hamilton glass syringes. The total flow rate was 0.5 mL/min, with LNP:cargo solutions mixing ratio 1:3. We used a passive herringbone mixer chip (Darwin Microfluidics) and the sample collection was started after the first 15 s. Following the microfluidic formulation, the LNP samples were dialyzed overnight at RT against DPBS++ (Gibco™, Thermo Fisher Scientific) in Slide-A-lyzer™ Mini dialysis tubes (Thermo Fisher Scientific; 3.5 kDa cut-off) to remove the ethanol and reach neutral pH. pH was checked with pH paper.

DLS was performed immediately after the dialysis using a Malvern Zetasizer Nano series (Nano - ZS) instrument. LNPs were diluted 1 uL in 499 uL DPBS++ (Gibco™ ) and measured at 25°C in low volume cuvettes. Cargo loading was evaluated with Quant-it™ RiboGreen RNA assay kit (R11490, Invitrogen) using the manufacturer protocol. For RiboGreen the LNP samples were diluted 1:20 with the TE buffer from the kit and the measurement was conducted for both intact and lysed LNPs (lysis with 2% (v/v) Triton X-100 in TE). Cargo loading efficiency (%) was calculated using the formula: $(c_{\text{after lysis}} - c_{\text{before lysis}})/c_{\text{after lysis}} * 100\%$. LNPs were stored at room temperature until the microscopy experiments were performed.

**Preparation of exosomes.** Cell culture and isolation of extracellular vesicles. EVs were prepared from HEK293-FS suspension cells (ThermoFisher), cbMSC (immortalized human cord blood-derived mesenchymal stromal cells, ATCC PCS-500-010), BJ-5ta (immortalized human fibroblast ATCC CRL-4001), and THP-1 (human monocytic cells). Cell lines were cultured in the following media: cbMSCs were cultured in MEM-α Modification Medium (containing L-glutamine; Thermo Fisher Scientific) supplemented with 5 ng/ml of bFGF (Sigma, F0291). BJ-5ta fibroblast cells were cultured with 4:1 mixture of Dulbecco's medium (containing 4 mM L-glutamine, 4.5 g/L glucose and 1.5 g/L sodium bicarbonate) and Medium 199 (0.01 mg/ml Hygromycin B/10687010, Thermo Fisher), HEK293-FS were cultured in FreeStyle 293 Expression Medium (ThermoFisher Scientific) in 125 mL polycarbonate Erlenmeyer flasks (Corning) in a shaking incubator (Infors HT Minitron) according to the manufacturer's instructions. THP-1 cells were cultured in RPMI-1640 medium (containing Glutamax-I and 25 mM HEPES, Invitrogen). Unless indicated otherwise, all cells were supplemented with 10% FBS (Invitrogen), 1X Antibiotic-Antimycotic (Anti-Anti) (Thermo Fisher Scientific). All cell lines were grown at 37°C, 5% CO2 in a humidified atmosphere and regularly tested for the presence of mycoplasma. For EV harvesting, cell culture-derived conditioned media (CM) was changed to OptiMem (Invitrogen) 48 h before harvest of conditioned media as described before (Nordin et al., 2019). Unless indicated otherwise, all conditioned media samples were directly subjected to a low-speed centrifugation step at 500 xg for 5 min followed by a 2,000 xg spin for 10 min to remove larger particles and cell debris. Pre-cleared cell culture supernatant was subsequently filtered through 0.22 µm bottle top vacuum filters (Corning, cellulose acetate, low protein binding) to remove any larger particles. EVs were isolated by tangential flow filtration (TFF). For the TFF EV isolation, pre-cleared CM was concentrated via TFF by using the KR2i TFF system (SpectrumLabs) equipped with modified polyethersulfone (mPES) hollow fiber filters with 300 kDa membrane pore size (MidiKros, 370 cm2 surface area, SpectrumLabs) at a flow rate of 100 mL/min (transmembrane pressure at 3.0 psi and shear rate at 3700 sec−1) as described previously [2](Nordin et al., 2019). Amicon Ultra-0.5 10 kDa MWCO spin-filters (Millipore) were used to concentrate the sample to a final volume of 100 µL. The sample was then loaded on a qEV column (Izon Science) and the EV fractions were collected according to the manufacturer's instructions.

**Preparation of virus-like particles.** Hek293T cells were cultured in DMEM supplemented with 10% FCS and tested mycoplasma-free. To produce pseudotyped non-fluorescent VLPs, cells were seeded at a confluency of ~70% in T75 flasks and co-transfected 6 hours later with 8.2µg of the lentiviral packaging vector psPAX2 (gift from Didier Trono (Addgene plasmid # 12260) and 10µg of the respective viral surface protein using jetOptimus© (Polyplus) according to the suppliers' recommendations. Human codon usage optimized plasmids encoding for SARS-CoV-2 spike variants delta/beta and Ebola GP were kindly provided by Benjamin Murrell and Jochen Bodem, respectively. After 12 hours media was exchanged and VLPs harvested twice after 24 hours, respectively. Enrichment of VLPs was performed using Lenti-X™ Concentrator (Takara) according to the protocol provided by the company. Alternatively, to increase the purity, supernatants with VLPs were filtered through 0.45µm PES filters and applied to ultracentrifugation using a 40% sucrose cushion.

To produce fluorescent VLPs, Hek293T cells were co-transfected using Lipofectamine 3000 and 15µg of the delta-spike expression plasmid, 7.5µg DNA encoding for HIV Vpr-GFP (NIH HIV Reagent Program, Division of AIDS, NIAID, NIH: pEGFP-Vpr, ARP-11386, contributed by Dr. Warner C. Greene), and 7.5µg encoding for a lentiviral packaging plasmid (psPAX2 was a gift from Didier Trono - Addgene plasmid # 12260). Media was exchanged after 12 hours and VLPs harvested after 24 and 48 hours and enriched fiftyfold using LentiX concentrator according to the protocol provided by the manufacturer (Takara). To generate VLPs with different spike-cleavage patterns, cells were kept in the presence of 50µM furin-inhibitor (Decanoyl-RVKR-CMK; Tocris: 3501) after the medium change.

**Single particle profiling measurements and analysis.** Single Particle Profiling was performed using the setup for fluorescence correlation spectroscopy (FCS) at a Zeiss LSM 780 microscope. A 488 nm argon ion laser was used for TopFluor Cholesterol , FastDiO, GFP and NR12S and a 633 nm He−Ne laser was used for ASR PE. A 40×1.2 NA water immersion objective was used to focus the light. The laser power was set to 0.1−0.5% of the total laser power that corresponds to 2−10 µW. The emission detection windows were set as 490 – 560 for TF Chol and 650 – 700 for ASR PE. Emission from NR12S was recorded simultaneously in both channels. 40 intensity fluctuation traces 15 seconds long were acquired for every sample.

Traces and curves were then analysed using the homemade python program. The source code as well as the standalone distributions for Windows and Mac are available at the Github: https://github.com/taras-sych/Single-particle-profiler/tree/Multiple-files/Standalone%20distributions

Briefly, individual peaks from the traces were identified and intensities for these individual peaks were extracted with further calculation of GP if applicable. Furthermore, based on these values, dot plots or density plots were constructed. The description and guide to the program is available as a video tutorial at https://www.youtube.com/watch?v=NAwmWDSHEV8

Moreover, diffusion analysis was also performed using the same homemade software. Curves were fitted with the following three-dimensional diffusion:

$$G(\tau) = \frac{1}{N}\left(1 + \frac{\tau}{\tau_D}\right)^{-1}\left(1 + \frac{\tau}{AR^2\tau_D}\right)^{-\frac{1}{2}}$$

where N represents the number of fluorescent species within the beam's focal volume. Next, the diffusion coefficients were calculated as follows:

$$D = \frac{\omega^2}{8\ln(2)\tau_D}$$

where *w* corresponds to the full width of half-maximum of the point spread function, $\tau_D$ is the diffusion time, and *D* is the diffusion coefficient.

**Statistical analysis**

For every box plot the exact sample size is shown in brackets. The replicates come from repeated measurement of the same biological sample. The significance was determined in all cases by nonparametric one way ANOVA test. For all box plots: central line indicates the mean, box indicates 25-75 % and whiskers indicate min and max of the data. The significance is indicated by stars: *$p>0.05$ – n.s., $p<0.05$ - *, $p<0.01$ - **, $p<0.001$ - ***, $p < 0.0001$ - ****.*

**Reporting summary.** Further information on research design is available in the Nature Research Reporting Summary linked to this article.

### Supplementary Figures

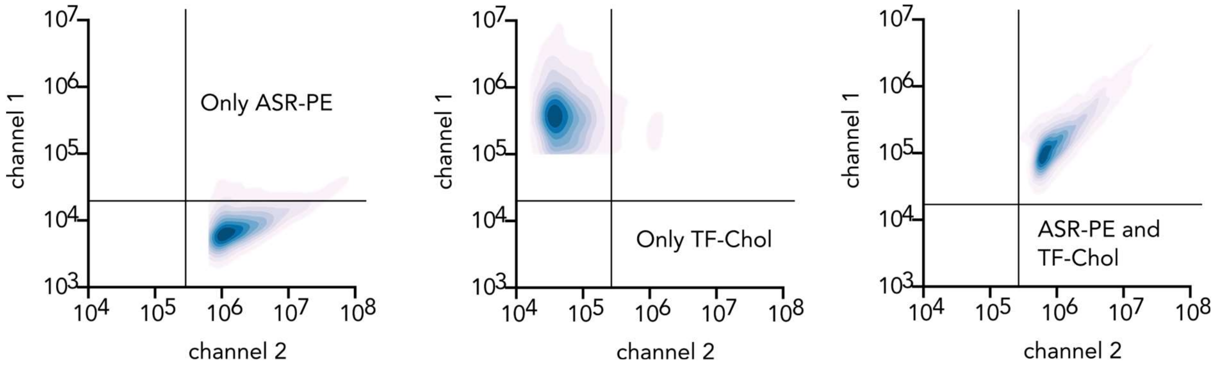


**Supplementary Figure 1.** Liposomes labelled with single fluorophore (TF-Chol: Topfluor Cholesterol; ASR-PE: Abberior STAR RED labelled phospholipid).

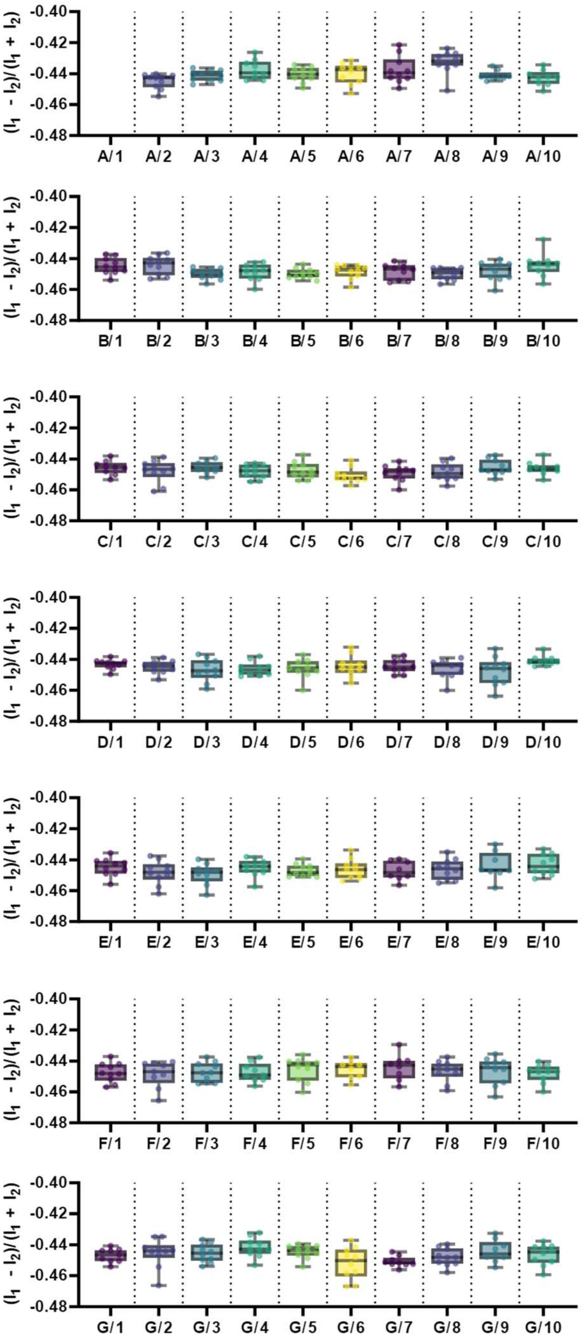


**Supplementary Figure 2.** High throughput analysis of multi-well plates with automated stage movement. The microscope stage moves once the chamber type and dimensions are introduced to the imaging program. Two channel recordings were done for 96-well plate with glass bottom. Corner wells could not be reached due to physical hindrances by the stage and the objective.

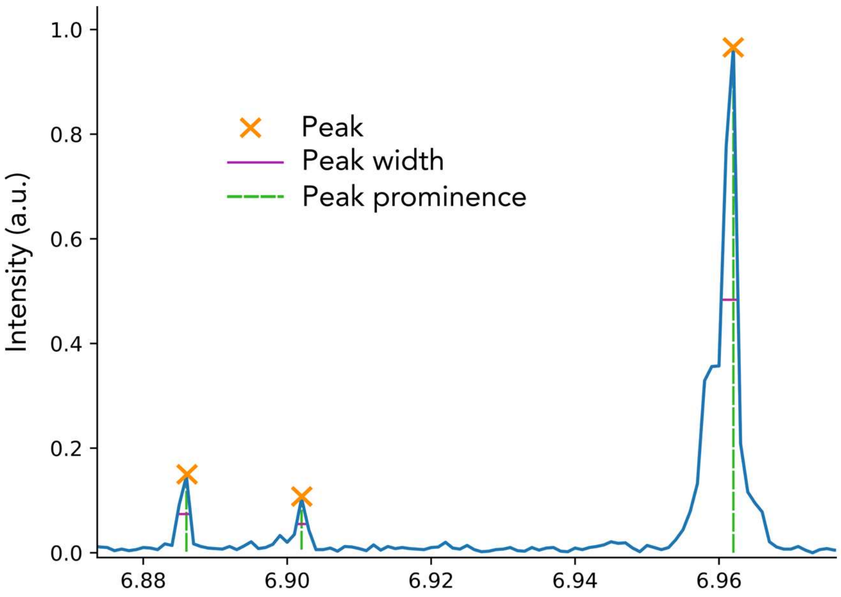


**Supplementary Figure 3.** Parameters that can be obtained for each peak.

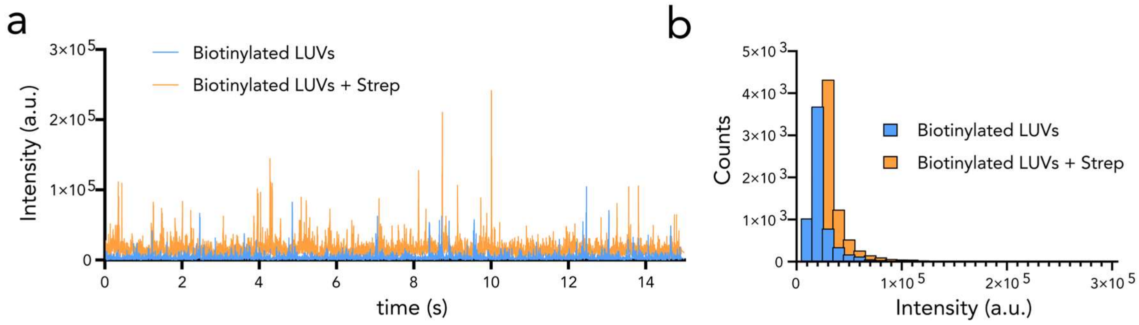


**Supplementary Figure 4.** Clusters of liposomes can be separated by brightness. a) Peaks caused by liposomes incorporated with fluorescent lipids and biotinylated lipids with and without streptavidin. Streptavidin causes aggregation, and the peak intensity increases accordingly; b) Intensity histograms before (blue) and after (prange) the introduction of streptavidin.

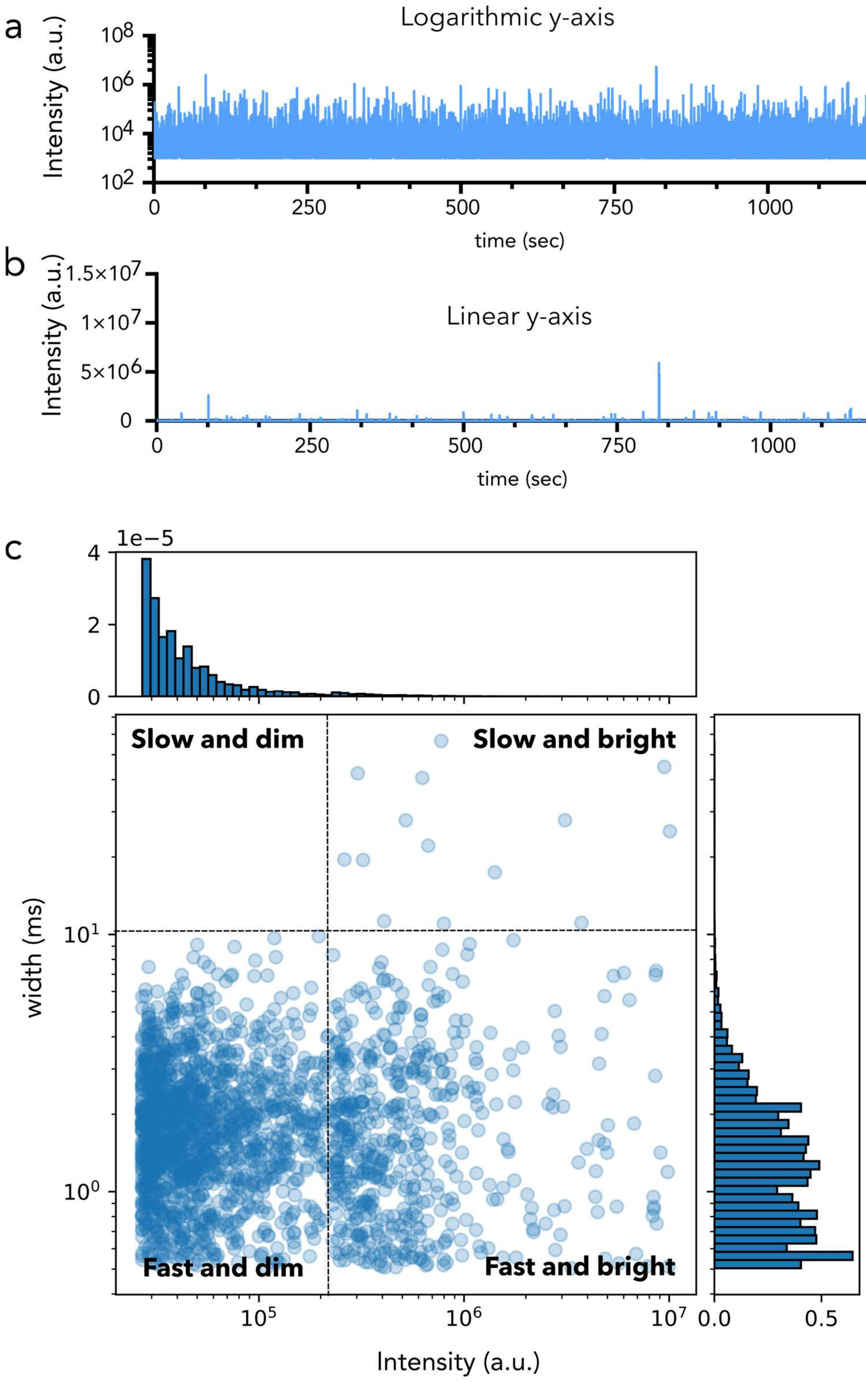


**Supplementary Figure 5.** Aggregated or big molecules can be distinguished clearly when peak width is combined with peak intensity. a, b) Peaks caused by an heterogenous liposome mixture (big and small liposomes) in linear vs logarithmic scales, respectively. c) Peak width vs intensity scatter plot shows slow/ bright particles compared to fast/bright and fast/dim particles.

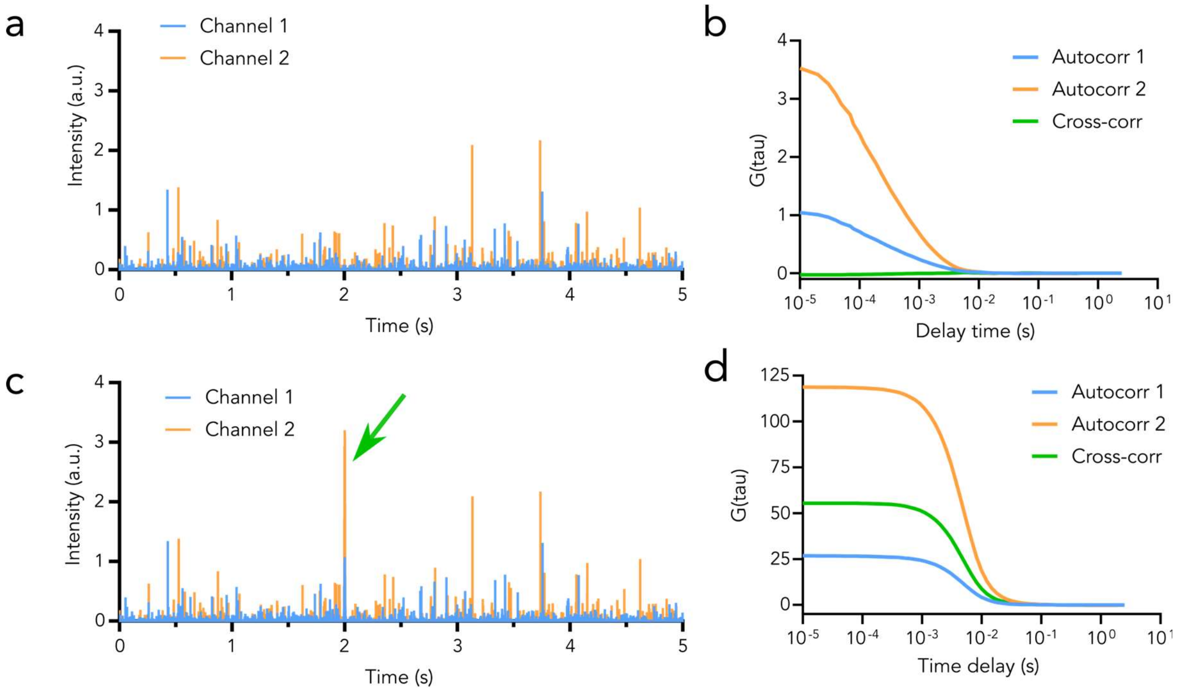


**Supplementary Figure 6.** Two colour fluorescence cross correlation spectroscopy cannot provide reliable information on number of co-occurrences in heterogeneous samples. a) Peaks caused by an homogenously distributed sample and b) cross-correlation curves showing no co-occurrences. c) the same sample with a single bright broad peak introduced artificially into both intensity traces at the same timepoint;d) the cross-correlation is now heavily biased by this single peak and shows 100% cross correlation. Data was generated by simulation via random-walk diffusion using home-made script in python. Script is freely available here: https://github.com/taras-sych/FCCS_simulation

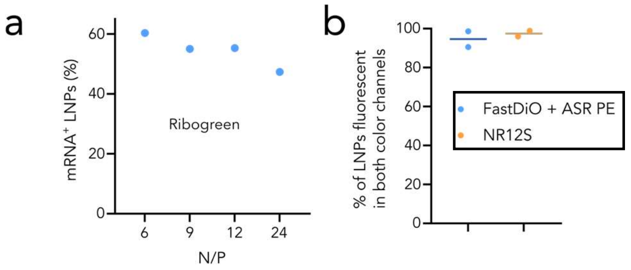


**Supplementary Figure 7.** LNP control experiments. a) RiboGreen analysis of LNPs with different N/P ratio. b) LNPs with green (Fast-DiO) and red lipid dyes (Abberior StarRed-PE) or a single dye with two emissions (NR12S) as a positive control for co-occurrence which show 100% co-occurrence.

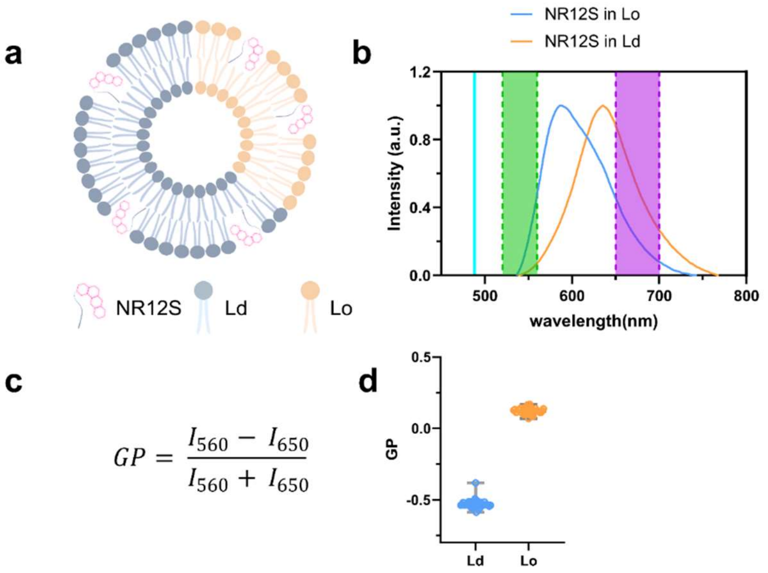


**Supplementary Figure 8. Measurement of membrane order using environmental sensitive dye. a.** Lipid bilayer consisting of two different domains, liquid ordered (Lo) and liquid disordered (Ld) labeled with NR12S; **b.** Fluorescence spectra of NR12S in Lo and Ld. Cyan line shows the excitation wavelength as well as green and magenta bands display the widths of the detection channels used for quantification in c; **c.** Quantification of the generalized polarization of NR12S spectra shown in b; **d.** GP of Lo and Ld.

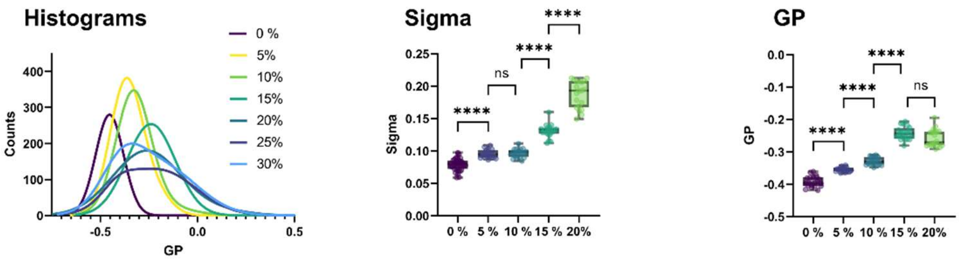


**Supplementary Figure 9. LUVs with different content of cholesterol.** Studies of mixtures of pure POPC with POPC/chol LUVs of different cholesterol content: GP histogram of mixtures, sigma of distribution, mean gp for single component distributions.

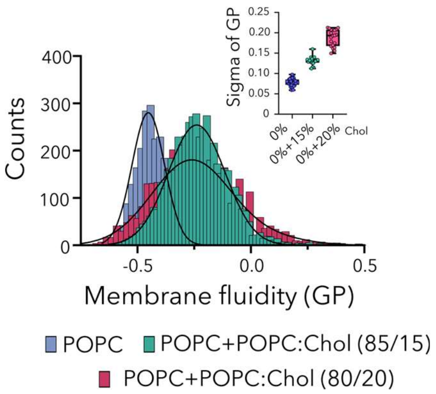


**Supplementary Figure 10. LUVs with different content of cholesterol.** Data for binary mixture of pure POPC liposomes and POPC/chol liposomes with different percentage of cholesterol. GP histograms show one population for 0% chol – 20% chol. Even where multiple populations cannot be resolved at low cholesterol mixtures, heterogeneity manifests in broadening of the distribution as reflected by the sigma values.

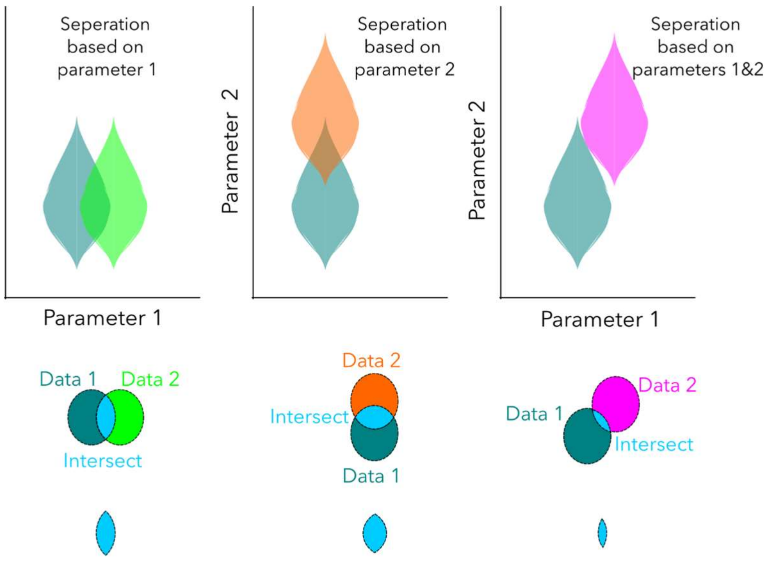


**Supplementary Figure 11.** Two parameters instead of one has higher separation power, with less overlap.

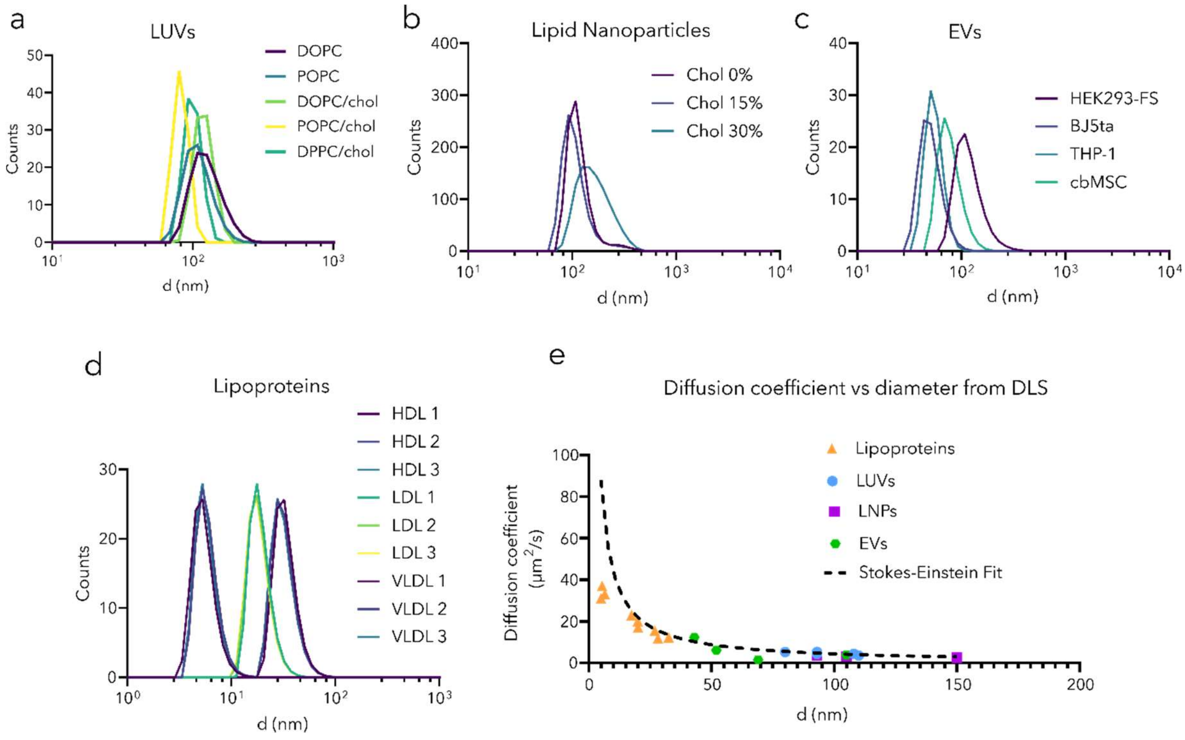


**Supplementary Figure 12.** Diffusion vs size from SPP measurements. a-d) DLS size measurements for liposomes, LNPs, EVs and lipoproteins. e) The relationship between the DLS size measurements and diffusion coefficient we obtained from SPP. Estimation is robust for all particles except HDLs.

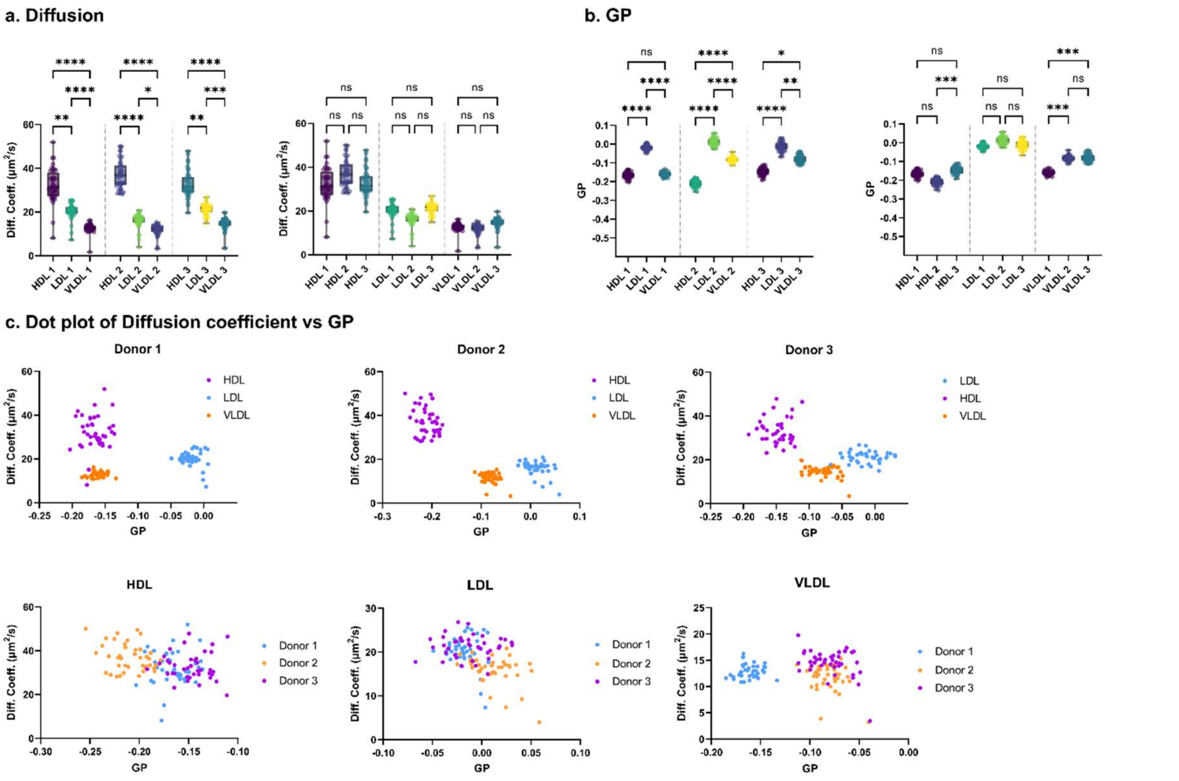


**Supplementary Figure 13. Lipoproteins from healthy individuals.** LPs were isolated from blood plasma of healthy individuals; **a.** Diffusion coefficient of lipoproteins; Left and right panel contain the same information, but on left panel it is sorted by donor and on right panel – by type of lipoprotein. **b.** GP of lipoproteins; Left and right panel contain the same information, but on left panel it is sorted by donor and on right panel – by type of lipoprotein. **c.** Dot plot of Diffusion Coefficient vs GP;

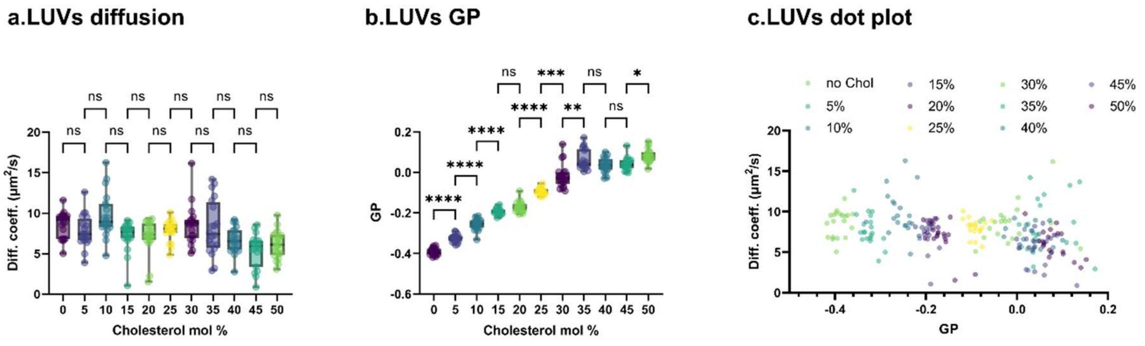


**Supplementary Figure 14. LUVs with different content of cholesterol.** LUVs that consist of POPC with different mol % of cholesterol were profiled. **a.** Diffusion coefficients of such LUVs; **b.** GP of such LUVs; **c.** Dot plot of Diffusion Coefficient vs GP.

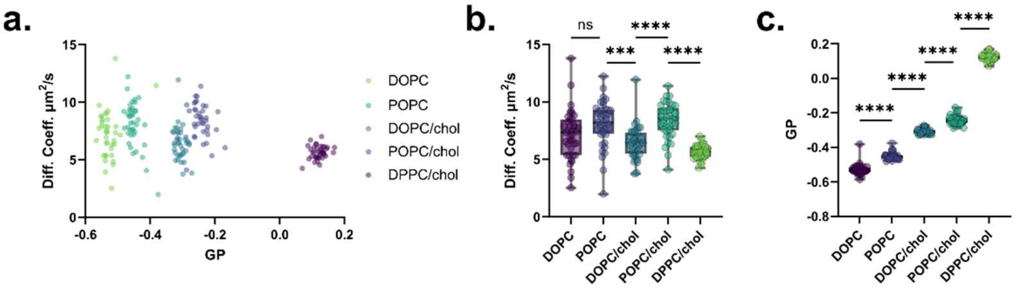


**Supplementary Figure 15. Profiling of LUVs.** LUVs were labeled by NR12S; **a.** Dot plot of Diffusion Coefficient vs GP; **b.** Diffusion coefficients of LUVs; **c.** GP of LUVs.

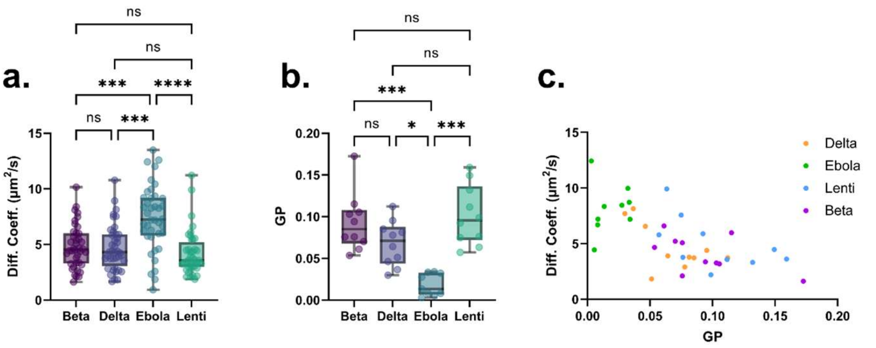


**Supplementary Figure 16. Virus-like particles (VLPs).** VLPs based on lentiviruses co transfected with SARS-COV-2 Spike glycoproteins (beta and delta mutations), ebola glycoprotein and lentiviruses without glycoproteins; **a.** VLPs diffusion; **b.** GP of VLPs; **c.** Dot plot of Diffusion Coefficient vs GP for VLPs;

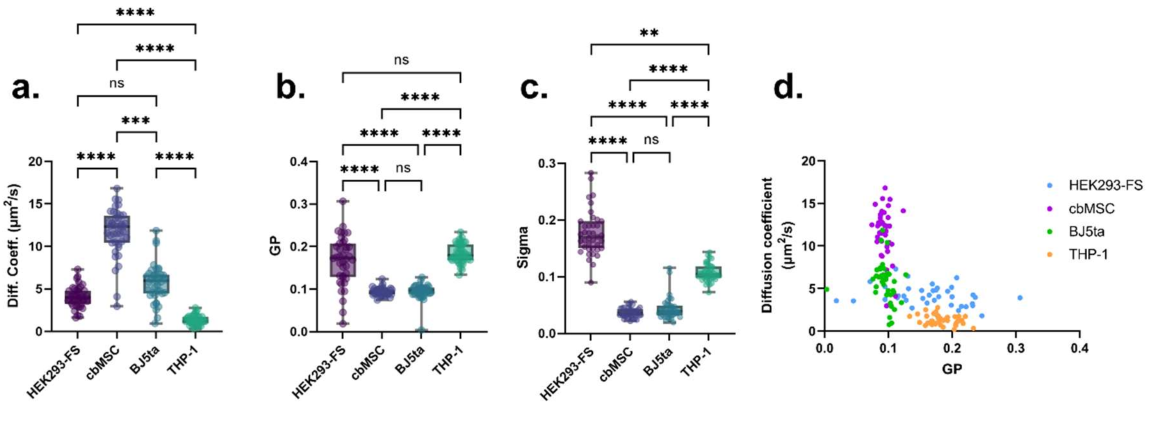


**Supplementary Figure 17. Exosomes isolated from different cells. a.** Exosome diffusion; **b.** GP of exosomes; **c.** Sigma of EVs; **d.** Dot plot of Diffusion Coefficient vs GP for exosomes.

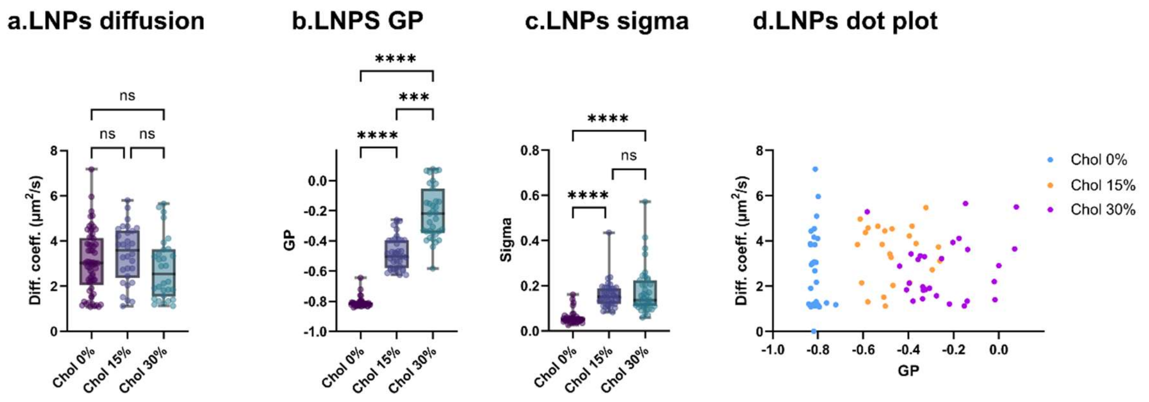


**Supplementary Figure 18. LNPs with different content of cholesterol. a.** Diffusion of lipid nanoparticles (LNPs); **b.** GP of LNPs; **c.** Sigma of LNPs **d.**Dot plot of LNPs;

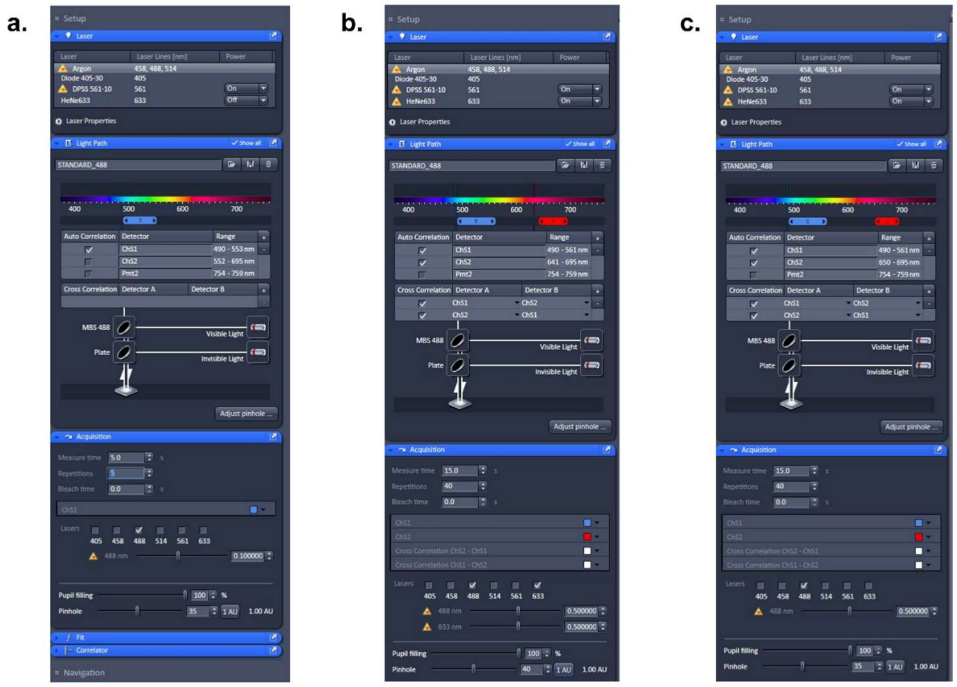

**Supplementary Figure 19. Settings for Fluorescence Correlation Spectroscopy measurements for SPP. a.** Settings for calibration with Alexa 488 in water. Excitation laser: 488 nm, detection window: 490-553 nm, 5s measurement time with 5 repetitions, laser power at 0.1%; **b.** Settings for two-color profiling with TF-chol and ASR-PE. Excitation lasers: 488 nm for TF-Chol and 633 nm for ASR-PE, detection windows: 490-561 nm for TF-Chol and 641-695 nm for ASR PE, 15s measurement time with 40 repetitions, laser powers at 0.5% **c.** Settings for biophysical profiling with ratiometric lipophilic probe NR12S. Excitation lasers: 488 nm, detection windows: 490-561 nm for green part of NR12S spectrum and 650-695 nm for red part of NR12S spectrum, 15s measurement time with 40 repetitions, laser power at 0.5%.
